# Genome Sequence of a Marine Threespine Stickleback (*Gasterosteus aculeatus*) from Rabbit Slough in the Cook Inlet

**DOI:** 10.1101/2025.02.06.636934

**Authors:** Eric H. Au, Seth Weaver, Anushka Katikaneni, Julia I. Wucherpfennig, Yanting Luo, Riley J. Mangan, Matthew A. Wund, Michael A. Bell, Craig B. Lowe

## Abstract

The Threespine Stickleback, *Gasterosteus aculeatus*, is an emerging model system for understanding the genomic basis of vertebrate adaptation. A strength of the system is that marine populations have repeatedly colonized freshwater environments, serving as natural biological replicates. These replicates have enabled researchers to efficiently identify phenotypes and genotypes under selection during this transition. While this repeated adaptation to freshwater has occurred throughout the northern hemisphere, the Cook Inlet in south-central Alaska has been an area of focus. The freshwater lakes in this area are being studied extensively and there is a high-quality freshwater reference assembly from a population in the region, Bear Paw Lake. Using a freshwater reference assembly is a potential limitation because genomic segments are repeatedly lost during freshwater adaptation. This scenario results in some of the key regions associated with marine-freshwater divergence being absent from freshwater genomes, and therefore absent from the reference assemblies. It may also be that isolated freshwater populations are more genetically diverged, potentially increasing reference biases. Here we present a highly-continuous marine assembly from Rabbit Slough in the Cook Inlet. All contigs are from long-read sequencing and have been ordered and oriented with Hi-C. The contigs are anchored to chromosomes and form a 454 Mbp assembly with an N50 of 1.3 Mbp, an L50 of 95, and a BUSCO score over 97%. The organization of the chromosomes in this marine individual is similar to existing freshwater assemblies, but with important structural differences, including the 3 previously known inversions that repeatedly separate marine and freshwater ecotypes. We anticipate that this high-quality marine assembly will more accurately reflect the ancestral population that founded the freshwater lakes in the area and will more closely match most other populations from around the world. This marine assembly, which includes the repeatedly deleted segments and offers a closer reference sequence for most populations, will enable more comprehensive and accurate computational and functional genomic investigations of Threespine Stickleback evolution.

## Introduction

Threespine Stickleback is an emerging model organism for studying the molecular basis of vertebrate adaptation (Peichel et al. 2001; Reid *et al*. 2021). A key strength of the system is that marine and sea-run (anadromous) populations have repeatedly colonized freshwater habitats. During these transitions from saltwater to freshwater environments, the same phenotypes have evolved in parallel. This repeated appearance of similar traits during similar transitions is consistent with the phenotypes being favored by natural selection in freshwater. These parallel adaptations to freshwater enable powerful analyses to uncover the genetic basis of vertebrate adaptations.

Examples of these repeatedly evolving traits include the number and size of armor plates (O’Brown et al. 2015; Indjeian *et al*. 2016), the loss of pelvic spines (Chan et al. 2010; Xie et al. 2019), and pigmentation differences (Miller et al. 2007). From these studies we have learned that the mutation selected during recent freshwater colonization events is often an allele from older freshwater populations that is present at low frequencies in marine/anadromous populations through gene flow (Colosimo *et al*. 2005; Jones *et al*. 2012; Lowe *et al*. 2018; Roberts Kingman *et al*. 2021). Alternatively, there are cases where the parallel adaptation is due to similar mutations repeatedly occurring (Chan *et al*. 2010; Xie *et al*. 2019), such as the repeated deletion of a pelvic enhancer for PITX1 leading to spine loss in freshwater populations (Chan *et al*. 2010; Xie *et al*. 2019).

While the scientific community has studied marine/anadromous Threespine Stickleback populations adapting to freshwater across the northern hemisphere (Klepaker 1993; Terekhanova *et al*. 2014; Kakioka *et al*. 2020; Aguirre *et al*. 2022; Venu *et al*. 2024), certain regions have made disproportionately large contributions to our understanding of the molecular and evolutionary mechanisms underlying vertebrate adaptation. This is due to both a wealth of colonization events and decades of sustained research efforts (Hagen and McPhail 1970; Schluter and McPhail 1992). One notable example is the Cook Inlet region of South-Central Alaska (Bell *et al*. 1985; Francis *et al*. 1986). The initial stickleback genome assembly came from Bear Paw Lake in this region (Jones et al. 2012), and the genetics of freshwater stickleback in the area are continuing to be studied intensively, including many lake populations founded recently by anadromous stickleback (Aguirre et al. 2022).

Currently, genomic studies using Threespine Stickleback rely on freshwater reference genomes. This is a potential limitation because the deletion of genomic segments is a prominent class of mutations when adapting to freshwater (Chan et al. 2010; Lowe et al. 2018; Thompson et al. 2018). This scenario results in some of the key regions associated with marine-freshwater divergence being absent from freshwater genomes, and therefore absent from the (freshwater) reference assemblies. Being absent in the reference assembly means that these key genomic segments are often not considered during computational or functional genomic screens that involve mapping sequence reads back to a reference genome. It may also be that isolated freshwater populations are more genetically diverged, potentially increasing reference biases. Together, these concerns could limit the efficiency, accuracy, and scope of comparative genomic analyses.

To address these challenges, we present a highly-continuous assembly of a marine/anadromous fish from Rabbit Slough (RABS), Alaska. It is likely that this assembly better reflects the genetics of the ancestral population that founded the extensively studied freshwater populations in the area. We anticipate that this highly-continuous marine/anadromous assembly from the Cook Inlet will enable more thorough computational and functional genomic screens in Threespine Stickleback.

## Materials and Methods

### Sample collection

For whole genome sequencing we used a female Threespine Stickleback fish that is the lab-reared offspring of fish originally collected from Rabbit Slough. The collection site is located where Rabbit Slough flows under the Parks Highway (61.5344N, 149.2677W). The RABS population is marine anadromous and associated with a fully-plated and pelvic-complete phenotype.

The Threespine Stickleback includes strictly marine, anadromous, and strictly freshwater populations (Bell and Foster 1994). The Rabbit Slough population is anadromous (Bell et al. 2016) and spends most of its life cycle in the ocean. We refer to it as “marine” for brevity and to emphasize that it spends most of its life cycle in marine waters, though it spends the first few months and last few weeks of life in fresh water.

### DNA extraction

We dissected brain and tail tissue. We immediately froze these tissue samples at −80° C. We later extracted high molecular weight genomic DNA from both tissues using the Qiagen MagAttact High-Molecular-Weight (HMW) DNA kit, per the manufacturer’s protocol.

### Whole genome sequencing

#### Pacific Biosciences

We sheared and size selected genomic DNA from the brain tissue to target an insert size of 20 kb. The Duke Sequencing and Genomic Technologies core facility performed library preparation and sequencing using the Sequel V3 chemistry, generating data across 12 SMRT Cells.

#### Oxford Nanopore

We also used genomic DNA from the same brain sample to prepare Ultra Long Oxford Nanopore Technology sequencing libraries using the Ligation Sequencing kit (SQK-LSK109) and sequenced the resulting libraries on 3 MinION flow cells.

#### 10x Linked-Reads

We used the same brain sample to generate genomic sequences with the linked reads technology from 10x Genomics. The Novogene Genome Sequencing Company used the 10x Chromium Controller to partition 0.6ng of fragmented high molecular weight genomic DNA and uniquely barcoded beads into GEMs (Gel bead-in-EMulsions) following the standard protocol of the Chromium Genome Reagent Kit User Guide (CG00022 RevA). Then they sequenced the resulting library on an Illumina HiSeq X.

### Hi-C

We generated Hi-C (Dudchenko *et al*. 2017) libraries from the flash-frozen tail tissue. We used the Arima Genomics chemistry, which utilizes four restriction enzymes for chromatin digestion. We labeled the fragment ends with biotin, ligated them by proximity, and purified, fragmented, and size selected for 300 base pairs. We enriched for ligation junctions and created Illumina-compatible sequencing libraries using the KAPA Hyper Prep with Illumina TruSeq adapters and indices. We sequenced the finished library on an Illumina HiSeq X.

We also used an existing Hi-C data set from a benthic freshwater Threespine Stickleback fish from Paxton Lake (Peichel *et al*. 2017) in our analyses, but not for genome assembly.

### Genome assembly

#### Contig construction and quality control

We performed the initial base call corrections, trimming of sequence adapters, and overlap contig consensus on an estimated 205x combined coverage of Pacific Biosciences and Oxford Nanopore Technologies data using CANU (Koren *et al*. 2017) for two rounds. The initial draft assembly contained multiple contigs representing the same genomic regions, which is commonly due to long stretches of elevated heterozygosity between the maternal and paternal chromosomes (Koren *et al*. 2011). We used Purged Haplotigs (Roach *et al*. 2018) on the initial draft assembly to identify and remove the redundant contigs. First, we mapped whole genome raw reads to the draft assembly using minimap2 (Li 2018). This produced a bimodal distribution of coverage representing both haploid and diploid levels of coverage. We used this information to flag contigs for manual inspection and potentially for removal, iteratively refining the assembly to arrive at a set of contigs with diploid coverage.

#### Building scaffolds

We started building scaffolds by mapping 10x Genomics Linked-Reads to the contigs. We generated alignment information with Long Ranger (Bishara *et al*. 2015), a modified BWA (Li and Durbin 2009) aligner that incorporates linked-read barcodes. We then used Scaff10x (github.com/wtsi-hpag/Scaff10X) to join contigs into scaffolds.

After initial assembly scaffolding, we used proximity-guided Hi-C libraries to both verify the assembly and improve scaffolding by comparing Hi-C heat map contacts (Durand *et al*. 2016). First, we performed Illumina adapter trimming with BBDuk (sourceforge.net/projects/bbmap) and Trim Galore (github.com/FelixKrueger/TrimGalore) (Martin 2011). We then aligned the paired-end reads independently with BWA (Li and Durbin 2009). We used software from Arima Genomics to output a sorted, mapping quality filtered, paired-end BAM file (github.com/ArimaGenomics/mapping_pipeline). We removed PCR duplicates using Picard (github.com/broadinstitute/picard) and converted the aligned BAM file to BED format with BEDTools (Quinlan and Hall 2010). We then used SALSA (Ghurye et al. 2017, 2019) and the Hi-C data to correct assembly errors and assemble additional scaffolds.

To order and orient scaffolds, we aligned our scaffolds to the freshwater stickleback assemblies, gasAcu1 (Jones et al. 2012) and GAculeatus_UGA_version5 (Nath et al. 2021). We used minimap2 (Li 2018) and MUMmer (Marçais et al. 2018) for the alignments and Ragoo (Alonge et al. 2019) for order and orientation. After this process, 18 scaffolds remained unordered. Using BLAST (Altschul *et al*. 1990), we determined that the 18 unordered scaffolds were repetitive elements already present in the assembly, sequence adapters, or contamination. We removed all 18 unordered scaffolds from the assembly.

To polish the genome assembly, we used the high-coverage Pacific Biosciences sequencing data along with software from Pacific Biosciences (i.e. pbmm2 and Quiver). These alignments served to close gaps, correct small insertions and deletions, and correct single base identities. We iterated this process until we did not accept any more modifications to the assembly.

In the final assembly polishing step, we used Long Ranger (Bishara et al. 2015) to align the 10x Genomics Linked-Reads to the current assembly and freebayes (Garrison and Marth 2012) to identify differences between the assembly and the reads. We chose freebayes because it can identify a diverse set of variants, including single-nucleotide polymorphisms (SNPs), small insertions and deletions (INDELs), multi-nucleotide polymorphisms (MNPs), and more complex events.

For quality control and validation, we visualized Hi-C contact maps with Juicer (Durand et al. 2016) and checked for completeness by identifying Actinopterygii BUSCOs (Benchmarking Universal SingleCopy Orthologs) (Simão et al. 2015; Seppey et al. 2019). We used the assemblyStats program from Gonomics to calculate assembly statistics (Au et al. 2023).

### Repetitive element annotation

We identified mobile elements and other repeats in the marine assembly using RepeatMasker v4.1.6 with the Vertebrates Dfam v3.8 library (Storer et al. 2021). During this analysis, we used the RMBlast v2.14.1 search engine with the slow search option. For comparison, we also analyzed the gasAcu1 (Jones et al. 2012) and GAculeatus_UGA_version5 (Nath et al. 2021) freshwater assemblies.

### Genome-wide alignments to transfer previous annotations

We generated genome-wide alignments between the freshwater gasAcu1 assembly (Jones et al. 2012) and the marine assembly we present here. We used LASTZ (Harris 2007) to produce local alignments between the assemblies. We then chained and filtered these alignments with UCSC Kent Utilities: axtChain, chain-Filter, chainAntiRepeat, chainMergeSort, chainSort, chainPreNet, chainNet, netSyntenic, netFilter, netChainSubset, and chainStitchId (Kent et al. 2003). We used axtChain to create chained pairwise alignments for each chromosome separately, and computed these chains with both the marine and freshwater genomes as the query and target. We filtered the chains with chainFilter and chainAntiRepeat, merged and sorted the files with chainMergeSort and chainSort, and only selected the best alignments at every region of the reference genome to achieve single-coverage syntenic alignments using chainPreNet, chainNet, netSyntenic, and netFilter. We then used netChainSubset and chainStitchId to generate the liftover chain file from the freshwater genome to the marine genome.

### Gene annotation

We produced RNA-seq data to guide gene annotation. To capture a large diversity of genes and isoforms, we conducted RNA-seq in the developing brain, which is a complex tissue comprised of many cell types (Braun *et al*. 2023). Isoform usage can change over time (Patowary *et al*. 2024), so we generated RNA-seq data from 4 different points in development. To capture both marine and freshwater transcriptomes, we performed this data collection in F1 hybrids.

We crossed fish from the Rabbit Slough (RABS) and Lake Matanuska (LMK) populations. At 1, 3, 4, and 5 days post-hatching we sacrificed a set of fish and isolated the brain tissue. We placed the tissue in TriZol and extracted RNA with the Direct-zol RNA Miniprep Plus kit. We used the Illumina TruSeq Stranded mRNA library prep kit to perform polyA selection on the mRNA, transcribe it into cDNA, ligate on Illumina sequence adapters, and perform PCR to enrich for molecules that can be sequenced. The libraries were then sequenced by Novogene on the Illumina HiSeqX platform. We utilized BRAKER3 (Gabriel *et al*. 2024) to predict protein-coding genes and provided the current freshwater assembly’s predicted proteins Nath *et al*. (2021); Dyer *et al*. (2025) and the RNA-seq data we generated to guide the model.

### Mapping diverse populations to the marine assembly

We identified a set of geographically varied whole-genome sequencing data sets from both marine and freshwater ecotypes. We used individuals from the following populations: Alaskan Marine (AKMA, SAMN02864913) and Alaskan Stream (AKST, SAMN02864935) from southern Alaska, marine and freshwater populations from the Little Campbell River in British Columbia (LITC_0_05, SAMN02781694; LITC_23_32, SAMN02781068), marine and freshwater populations from the Big River in California (BIGR_1_32, SAMN02781111; BIGR_52_54, SAMN02781687), and marine and freshwater populations from the River Tyne in Scotland (TYNE_1, SAMN02781690; TYNE_8, SAMN02781066) (Roberts Kingman *et al*. 2021). We mapped reads to each of the assemblies with BWA (Li and Durbin 2009) and identified the percentage of reads that could not be mapped with Samtools (Li and Durbin 2009).

## Results and Discussion

### Assembly

#### Completeness

We present a highly contiguous marine stickleback genome assembly (*Duke*_*GAcu*_1.0), with all contigs placed onto one of the 21 nuclear chromosomes or the mitochondrial chromosome. The total length of the marine assembly (≈ 454mb) is similar to previous freshwater assemblies, with slightly more (2% increase) bases than the original assembly, gasAcu1 (Jones *et al*. 2012). The more recent freshwater reference assembly (Nath *et al*. 2021) is slightly larger (≈468mb), due to the inclusion of the Y chromosome, which is not present in our marine assembly from a female fish (Table 1).

**Table 1.** Assembly Statistics.

| Assembly | Ecotype | Size (bp) | Contigs | Longest contig (kb) | N50 (kb) | L50 |
| --- | --- | --- | --- | --- | --- | --- |
| gasAcu1 | Freshwater | 446,635,014 | 16,957 | 698.2 | 83.2 | 1,459 |
| GAculeatus_UGA_version5 | Freshwater | 468,320,375 | 6,058 | 4,692.7 | 485.8 | 253 |
| GAculeatus_UGA_version5 (no chrY) | Freshwater | 452,484,949 | 6,010 | 4,692.7 | 480.7 | 246 |
| Duke_GAcu_1.0 | Marine | 454,257,695 | 737 | 5,528.6 | 1,287 | 95 |

To evaluate the completeness of the marine assembly, we analyzed the BUSCO (Simão *et al*. 2015) results. Out of 3640 BUSCOs (proteins based on Actinopterygii OrthoDBv9), the marine assembly matched 3495 completely and with a single-copy; 43 are complete and duplicated, 7 are fragmented, and 95 are either missing or un-mappable, for an overall score of 97.2 percent. This is similar to the reported score for the current freshwater assembly (96.7%) (Nath *et al*. 2021), and consistent with a near-complete assembly of the euchromatic regions.

#### Continuity

While the freshwater and marine assemblies exhibit comparable completeness, the marine assembly demonstrates substantial improvements across metrics of continuity. The marine assembly has 88% fewer contigs, a 62% reduction in L50, and a 165% improvement in N50 (Table 1). Together, these data demonstrate that the marine assembly contains a similar amount of genomic sequence to previous freshwater assemblies, but with greater continuity.

#### Mobile elements and other repeats

We identified a relatively low level of repetitive DNA content (3.79%). This value is consistent with our analysis of previously published freshwater stickleback assemblies: 3.21% for gasAcu1 and 3.45% for GAculeatus_UGA_version5 Table 2. This low level of reported repeat content is likely to be from a mixture of the Threespine Stickleback still being an emerging model organism that has had less repeat analysis than the human or mouse genomes, as well as the Threespine Stickleback having a compact genome with low repeat content (Reid et al. 2021).

**Table 2.** Interspersed repeats in marine *G. aculeatus*.

| Name | Number of Elements | Total Length (bp) | Percent of Assembly |
| --- | --- | --- | --- |
| **Retroelements** | 25,529 | 3,133,799 | 0.69 |
| SINEs | 6,769 | 585,644 | 0.13 |
| LINEs | 8,527 | 1,681,574 | 0.37 |
| L2/CR1/Rex | 2,494 | 843,107 | 0.19 |
| RTE/Bov-B | 414 | 48,951 | 0.01 |
| L1/CIN4 | 5,848 | 819,488 | 0.18 |
| LTR elements | 10,233 | 866,581 | 0.19 |
| Gypsy/DIRS1 | 222 | 104,938 | 0.02 |
| Retroviral | 9,825 | 749,472 | 0.16 |
| **DNA transposons** | 6,517 | 901,701 | 0.20 |
| hobo-Activator | 3,540 | 407,029 | 0.09 |
| Tc1-IS630-Pogo | 2,918 | 489,716 | 0.11 |
| MULE-MuDR | 1 | 45 | 0.00 |
| PiggyBac | 31 | 2,371 | 0.00 |
| Tourist/Harbinger | 1 | 35 | 0.00 |
| **Rolling-circles** | 1,330 | 85,664 | 0.02 |
| **Unclassified** | 588 | 35,500 | 0.01 |
| **Total interspersed** |  | 4,071,000 | 0.90 |
| **Small RNA** | 9,362 | 771,590 | 0.17 |
| **Satellites** | 2,885 | 236,908 | 0.05 |
| **Simple repeats** | 257,902 | 10,772,290 | 2.37 |
| **Low complexity** | 26,103 | 1,341,067 | 0.29 |
| **Bases masked** |  | 17,215,130 | 3.79 |

#### Genes

We identified 21,494 predicted protein-coding genes using both RNA-seq data from the developing brain of F1 hybrids (RABS x LMK) as well as protein sequences from the most recent freshwater assembly (Nath et al. 2021; Dyer et al. 2025). This number is within the range (20,787 to 22,376) that has previously been reported by Ensembl for different freshwater genome assemblies (Jones et al. 2012; Nath et al. 2021; Dyer et al. 2025).

### Structural similarities and differences

While large-scale karyotypic differences have been characterized between stickleback species (Chen and Reisman 1970; Kitano *et al*. 2009; Urton et al. 2011), the high-level chromosomal organization is very similar between the marine and freshwater assemblies for Threespine Stickleback. We relied heavily on Hi-C contacts to assemble the marine genome from Rabbit Slough and it is interesting to note that Hi-C contacts from a freshwater fish (Peichel et al. 2017) provide largely the same signal across the genome (Figure 1). This result is consistent with the reproductive compatibility between freshwater and marine ecotypes (Reid et al. 2021; Schluter *et al*. 2021; Wucherpfennig et al. 2022), and with the freshwater populations in this region being founded after the last glacial maximum, only 21,000 years ago (Bell and Foster 1994; Lambeck et al. 2014).

**Figure 1.**
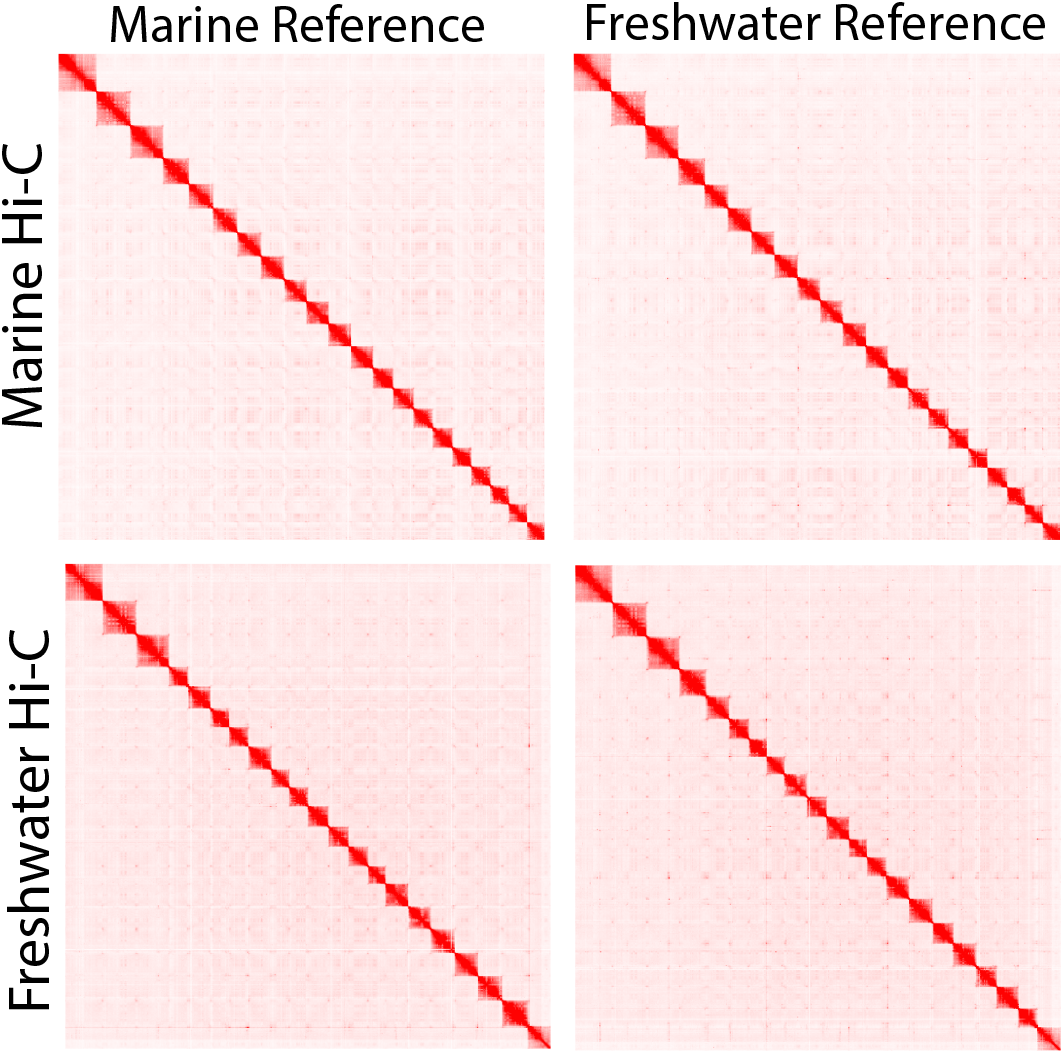
Marine and freshwater assemblies have nearly identical high-level chromosomal organization as evidenced by Hi-C contact maps from marine and freshwater (Peichel *et al*. 2017) individuals giving similar signals when mapped to marine or freshwater (Nath *et al*. 2021) assemblies.

**Figure 2.**
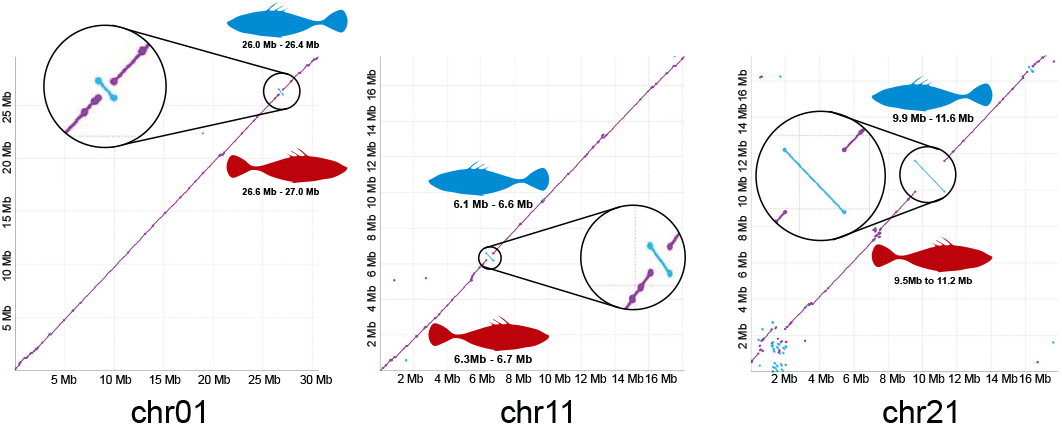
Discovery of known marine-freshwater inversions. Previous research identified 3 inversions that are repeatedly divergent between marine and freshwater populations (Jones et al. 2012). We discover all three of these inversions when comparing the marine genome we present here to the freshwater reference genome (Nath et al. 2021). These inversions serve a role similar to positive controls, which increases our confidence in the order and orientation of the genomic segments in the assembly.

While the high-level chromosomal arrangements between the marine and freshwater assemblies are similar, structural differences are present. Notably, our marine assembly captures 3 inversions on chrI/chr01, chrXI/chr11, and chrXXI/chr21 that were previously identified to repeatedly segregate between marine and freshwater individuals (Jones et al. 2012). Thus, while the overall genome structure is largely consistent between ecotypes, the marine assembly captures structural variants that are key components of ecotype divergence.

### Mapping efficiency

We investigated if the marine assembly is more representative of many stickleback populations, and therefore enables more comprehensive genomic analyses of Threespine Stickleback from around the world. We identified a collection of short-read whole-genome sequencing data sets from varied geographic locations. This consists of matched marine and freshwater ecotypes from Alaska, British Columbia, California, and Scotland (Roberts Kingman *et al*. 2021). Sequencing reads from all of these samples show better mapping to the marine assembly. A greater number of reads are mapped to the marine assembly, which should increase the accuracy and scope of genomic analyses Figure 3.

**Figure 3.**
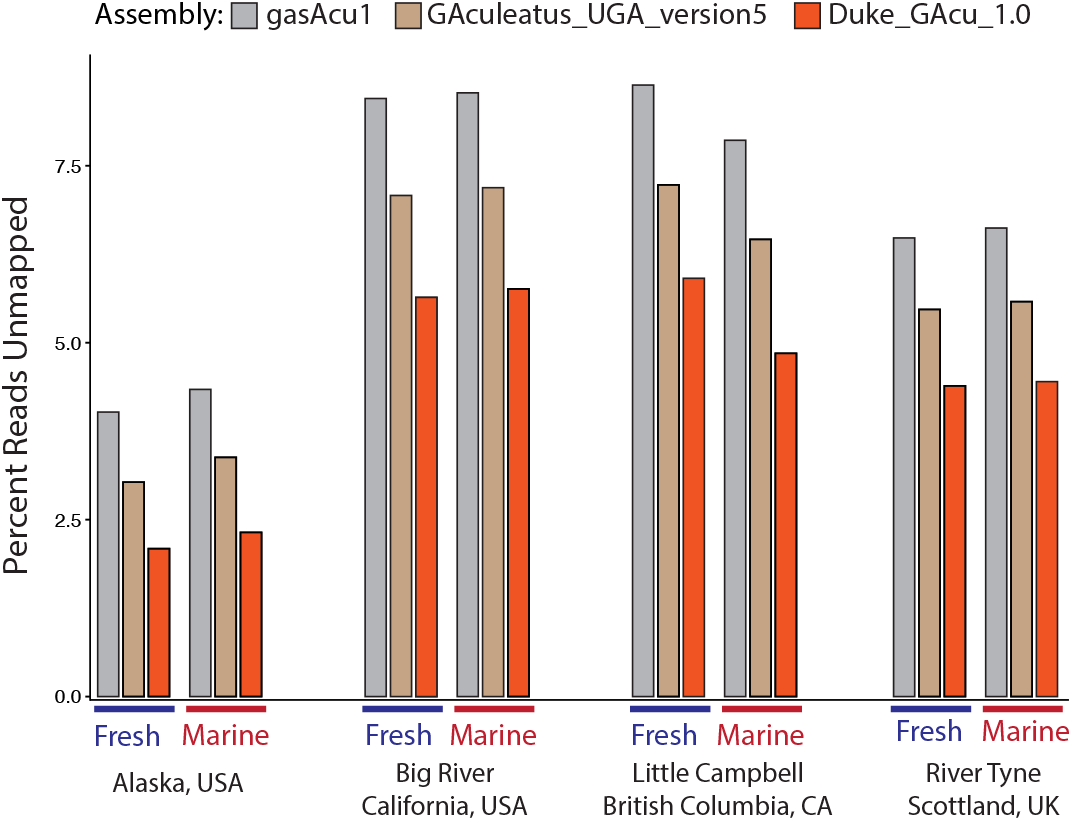
The marine assembly is a closer match to many populations from around the world, which allows for more reads to be mapped to the assembly and therefore a more comprehensive understanding of Threespine Stickleback genetics. We mapped reads from a variety of marine and freshwater populations to the two freshwater genome assemblies and the marine assembly we present here. We consistently observe a reduction in the percent of unmapped reads when using the marine genome, irregardless of the ecotype or geographic location.

## Discussion

Historically, it was common for a species to have a single reference genome, and the genetic diversity of the species to be understood by mapping short sequencing reads to that single reference (1000 Genomes Project Consortium et al. 2015). Reference genomes continually improved until we achieved telomere-to-telomere assemblies that represent single uninterrupted haplotypes (Nurk *et al*. 2022). We are now in the age where we have multiple assemblies to better understand the genetic diversity within a species (He *et al*. 2023; Yang et al. 2023). These multiple assemblies help researchers understand what structural variation is present within the species, as well as provide researchers with a set of assembly choices so that they can map short sequencing reads to an assembly that is closely related to the sequenced individual. Having a genetically similar reference genome is advantageous because sticklebacks (Chan et al. 2010; Lowe et al. 2018), humans (Sherman et al. 2019), and likely many other vertebrates have a large number of genomic segments that are present in one population, but may be absent from another. Our current challenge is to not only use a single reference that is genetically similar to our sample of interest, but combine all available assemblies into a single representation of a species (Liao et al. 2023) when analyzing future genomic data sets.

This will minimize the chance of information loss when reads are not correctly mapped to the reference genome, and offer a unified data structure where analyses from different research groups can be easily compared.

Another future challenge for the stickleback community is to generate and share functional genomic data sets. While there are incredible advantages to the stickleback system, an advantage of other systems is a wealth of functional genomic data sets (Mouse ENCODE Consortium *et al*. 2012; Roadmap Epigenomics Consortium *et al*. 2015; ENCODE Project Consortium *et al*. 2020; Boix *et al*. 2021). An advantage of the stickleback system is a relatively compact genome, approximately ^1^ the size of the human genome, which makes the generation of these data sets cost-effective. Groups are already generating these data sets on a small scale and finding them effective (Okude *et al*. 2024). We propose that a consortium of labs working together to generate functional genomic data for a large number of cell types across a diversity of populations would leverage strengths of the stickleback system and be cost-effective with current technologies.

## Data availability

We uploaded the sequencing data that we generated for this manuscript to NCBI. The data sets are freely available under BioProject number PRJNA1198983. Code written for this manuscript is freely available at https://github.com/vertgenlab/gonomics.

## Acknowledgments

We thank the Duke Sequencing Core for assistance with this project. The research reported here is covered by permits and protocols, including Alaska Department of Fish and Game permit SF-23-015, The College of New Jersey IACUC protocol 2208-001MW1, and Duke University IACUC protocol A166-21-08-24.

## Literature cited

1000 Genomes Project Consortium, Auton A, Brooks LD, Durbin RM, Garrison EP, Kang HM, Korbel JO, Marchini JL, McCarthy S, McVean GA et al. 2015. A global reference for human genetic variation. Nature. 526:68–74.

Aguirre WE, Reid K, Rivera J, Heins DC, Veeramah KR, Bell MA. 2022. Freshwater colonization, adaptation, and genomic divergence in threespine stickleback. Integr. Comp. Biol.. 62:388–405.

Alonge M, Soyk S, Ramakrishnan S, Wang X, Goodwin S, Sedlazeck FJ, Lippman ZB, Schatz MC. 2019. RaGOO: fast and accurate reference-guided scaffolding of draft genomes. Genome Biol. 20:224.

Altschul SF, Gish W, Miller W, Myers EW, Lipman DJ. 1990. Basic local alignment search tool. Journal of molecular biology. 215:403–410.

Au EH, Fauci C, Luo Y, Mangan RJ, Snellings DA, Shoben CR, Weaver S, Simpson SK, Lowe CB. 2023. Gonomics: uniting high performance and readability for genomics with go. Bioinformatics. 39.

Bell MA, Foster SA. 1994. The evolutionary biology of the threespine stickleback. Oxford University Press.

Bell MA, Francis RC, Havens AC. 1985. Pelvic reduction and its directional asymmetry in threespine sticklebacks from the cook inlet region, alaska. Copeia. 1985:437–444.

Bell MA, Heins DC, Wund MA, von Hippel FA, Massengill R, Dunker KJ, Bristow GA, Aguirre WE. 2016. Reintroduction of threespine stickleback into cheney and scout lakes, alaska. Evolutionary Ecology Research. 17:157–178.

Bishara A, Liu Y, Weng Z, Kashef-Haghighi D, Newburger DE, West R, Sidow A, Batzoglou S. 2015. Read clouds uncover variation in complex regions of the human genome. Genome Res.. 25:1570–1580.

Boix CA, James BT, Park YP, Meuleman W, Kellis M. 2021. Regulatory genomic circuitry of human disease loci by integrative epigenomics. Nature. 590:300–307.

Braun E, Danan-Gotthold M, Borm LE, Lee KW, Vinsland E, Lonnerberg P, Hu L, Li X, He X, Andrusivova Z et al. 2023. Comprehensive cell atlas of the first-trimester developing human brain. Science. 382:eadf1226.

Chan YF, Marks ME, Jones FC, Villarreal, Jr G, Shapiro MD, Brady SD, Southwick AM, Absher DM, Grimwood J, Schmutz J et al. 2010. Adaptive evolution of pelvic reduction in sticklebacks by recurrent deletion of a pitx1 enhancer. Science. 327:302–305.

Chen TR, Reisman HM. 1970. A comparative chromosome study of the north american species of sticklebacks (teleostei: Gasterosteidae). Cytogenetics. 9:321–332.

Colosimo PF, Hosemann KE, Balabhadra S, Villarreal, Jr G, Dickson M, Grimwood J, Schmutz J, Myers RM, Schluter D, Kingsley DM. 2005. Widespread parallel evolution in sticklebacks by repeated fixation of ectodysplasin alleles. Science. 307:1928–1933.

Dudchenko O, Batra SS, Omer AD, Nyquist SK, Hoeger M, Durand NC, Shamim MS, Machol I, Lander ES, Aiden AP et al. 2017. De novo assembly of the aedes aegypti genome using Hi-C yields chromosome-length scaffolds. Science. 356:92–95.

Durand NC, Robinson JT, Shamim MS, Machol I, Mesirov JP, Lander ES, Aiden EL. 2016. Juicebox provides a visualization system for Hi-C contact maps with unlimited zoom. Cell Syst.. 3:99–101.

Dyer SC, Austine-Orimoloye O, Azov AG, Barba M, Barnes I, Barrera-Enriquez VP, Becker A, Bennett R, Beracochea M, Berry A et al. 2025. Ensembl 2025. Nucleic Acids Res. 53:D948–D957.

ENCODE Project Consortium, Moore JE, Purcaro MJ, Pratt HE, Epstein CB, Shoresh N, Adrian J, Kawli T, Davis CA, Dobin A et al. 2020. Expanded encyclopaedias of DNA elements in the human and mouse genomes. Nature. 583:699–710.

Francis RC, Baumgartner JV, Havens AC, Bell MA. 1986. Historical and ecological sources of variation among lake populations of threespine sticklebacks, gasterosteus aculeatus, near cook inlet, alaska. Canadian Journal of Zoology. 64:2257–2265.

Gabriel L, na T, Hoff KJ, Ebel M, Lomsadze A, Borodovsky M, Stanke M. 2024. BRAKER3: Fully automated genome annotation using RNA-seq and protein evidence with GeneMark-ETP, AUGUSTUS, and TSEBRA. Genome Res. 34:769–777.

Garrison E, Marth G. 2012. Haplotype-based variant detection from short-read sequencing.

Ghurye J, Pop M, Koren S, Bickhart D, Chin CS. 2017. Scaffolding of long read assemblies using long range contact information. BMC Genomics. 18:527.

Ghurye J, Rhie A, Walenz BP, Schmitt A, Selvaraj S, Pop M, Phillippy AM, Koren S. 2019. Integrating Hi-C links with assembly graphs for chromosome-scale assembly. PLoS Comput. Biol.. 15:e1007273.

Hagen DW, McPhail JD. 1970. The species problem within gasterosteus aculeatus on the pacific coast of north america. Journal of the Fisheries Research Board of Canada. 27:147–155.

Harris RS. 2007. Improved pairwise alignment of genomic DNA. Ph.D. thesis. The Pennsylvania State University.

He Y, Chu Y, Guo S, Hu J, Li R, Zheng Y, Ma X, Du Z, Zhao L, Yu W et al. 2023. T2T-YAO: A telomere-to-telomere assembled diploid reference genome for han chinese. Genomics Proteomics Bioinformatics. 21:1085–1100.

Indjeian VB, Kingman GA, Jones FC, Guenther CA, Grimwood J, Schmutz J, Myers RM, Kingsley DM. 2016. Evolving new skeletal traits by cis-regulatory changes in bone morphogenetic proteins. Cell. 164:45–56.

Jones FC, Grabherr MG, Chan YF, Russell P, Mauceli E, Johnson J, Swofford R, Pirun M, Zody MC, White S et al. 2012. The genomic basis of adaptive evolution in threespine sticklebacks. Nature. 484:55–61.

Kakioka R, Mori S, Kokita T, Hosoki TK, Nagano AJ, Ishikawa A, Kume M, Toyoda A, Kitano J. 2020. Multiple waves of freshwater colonization of the three-spined stickleback in the japanese archipelago. BMC Evol. Biol.. 20:143.

Kent WJ, Baertsch R, Hinrichs A, Miller W, Haussler D. 2003. Evolution’s cauldron: duplication, deletion, and rearrangement in the mouse and human genomes. Proc. Natl. Acad. Sci. U. S. A.. 100:11484–11489.

Kitano J, Ross JA, Mori S, Kume M, Jones FC, Chan YF, Absher DM, Grimwood J, Schmutz J, Myers RM et al. 2009. A role for a neo-sex chromosome in stickleback speciation. Nature. 461:1079–1083.

Klepaker T. 1993. Morphological changes in a marine population of threespined stickleback, gasterosteus aculeatus, recently isolated in fresh water. Canadian Journal of Zoology. 71:1251–1258.

Koren S, Treangen TJ, Pop M. 2011. Bambus 2: scaffolding metagenomes. Bioinformatics. 27:2964–2971.

Koren S, Walenz BP, Berlin K, Miller JR, Bergman NH, Phillippy AM. 2017. Canu: scalable and accurate long-read assembly via adaptive k-mer weighting and repeat separation. Genome Res.. 27:722–736.

Lambeck K, Rouby H, Purcell A, Sun Y, Sambridge M. 2014. Sea level and global ice volumes from the last glacial maximum to the holocene. Proc. Natl. Acad. Sci. U. S. A.. 111:15296–15303.

Li H. 2018. Minimap2: pairwise alignment for nucleotide sequences. Bioinformatics. 34:3094–3100.

Li H, Durbin R. 2009. Fast and accurate short read alignment with Burrows-Wheeler transform. Bioinformatics. 25:1754–1760.

Liao WW, Asri M, Ebler J, Doerr D, Haukness M, Hickey G, Lu S, Lucas JK, Monlong J, Abel HJ et al. 2023. A draft human pangenome reference. Nature. 617:312–324.

Lowe CB, Sanchez-Luege N, Howes TR, Brady SD, Daugherty RR, Jones FC, Bell MA, Kingsley DM. 2018. Detecting differential copy number variation between groups of samples. Genome Res.. 28:256–265.

Marçais G, Delcher AL, Phillippy AM, Coston R, Salzberg SL, Zimin A. 2018. MUMmer4: A fast and versatile genome alignment system. PLoS Comput. Biol.. 14:e1005944.

Martin M. 2011. Cutadapt removes adapter sequences from highthroughput sequencing reads. EMBnet.journal. 17:10–12.

Miller CT, Beleza S, Pollen AA, Schluter D, Kittles RA, Shriver MD, Kingsley DM. 2007. cis-regulatory changes in kit ligand expression and parallel evolution of pigmentation in sticklebacks and humans. Cell. 131:1179–1189.

Mouse ENCODE Consortium, Stamatoyannopoulos JA, Snyder M, Hardison R, Ren B, Gingeras T, Gilbert DM, Groudine M, Bender M, Kaul R et al. 2012. An encyclopedia of mouse DNA elements (mouse ENCODE). Genome Biol.. 13:418.

Nath S, Shaw DE, White MA. 2021. Improved contiguity of the threespine stickleback genome using long-read sequencing. G3 (Bethesda). 11.

Nurk S, Koren S, Rhie A, Rautiainen M, Bzikadze AV, Mikheenko A, Vollger MR, Altemose N, Uralsky L, Gershman A et al. 2022. The complete sequence of a human genome. Science. 376:44–53.

O’Brown NM, Summers BR, Jones FC, Brady SD, Kingsley DM. 2015. A recurrent regulatory change underlying altered expression and wnt response of the stickleback armor plates gene EDA. Elife. 4:e05290.

Okude G, Yamasaki YY, Toyoda A, Mori S, Kitano J. 2024. Genome-wide analysis of histone modifications can contribute to the identification of candidate cis-regulatory regions in the threespine stickleback fish. BMC Genomics. 25:685.

Patowary A, Zhang P, Jops C, Vuong CK, Ge X, Hou K, Kim M, Gong N, Margolis M, Vo D et al. 2024. Developmental isoform diversity in the human neocortex informs neuropsychiatric risk mechanisms. Science. 384:eadh7688.

Peichel CL, Nereng KS, Ohgi KA, Cole BL, Colosimo PF, Buerkle CA, Schluter D, Kingsley DM. 2001. The genetic architecture of divergence between threespine stickleback species. Nature. 414:901–905.

Peichel CL, Sullivan ST, Liachko I, White MA. 2017. Improvement of the threespine stickleback genome using a Hi-C-based proximity-guided assembly. J. Hered.. 108:693–700.

Quinlan AR, Hall IM. 2010. BEDTools: a flexible suite of utilities for comparing genomic features. Bioinformatics. 26:841–842.

Reid K, Bell MA, Veeramah KR. 2021. Threespine stickleback: A model system for evolutionary genomics. Annu. Rev. Genomics Hum. Genet.. 22:357–383.

Roach MJ, Schmidt SA, Borneman AR. 2018. Purge haplotigs: allelic contig reassignment for third-gen diploid genome assemblies. BMC Bioinformatics. 19:460.

Roadmap Epigenomics Consortium, Kundaje A, Meuleman W, Ernst J, Bilenky M, Yen A, Heravi-Moussavi A, Kheradpour P, Zhang Z, Wang J et al. 2015. Integrative analysis of 111 reference human epigenomes. Nature. 518:317–330.

Roberts Kingman GA, Vyas DN, Jones FC, Brady SD, Chen HI, Reid K, Milhaven M, Bertino TS, Aguirre WE, Heins DC et al. 2021. Predicting future from past: The genomic basis of recurrent and rapid stickleback evolution. Sci. Adv.. 7:eabg5285.

Schluter D, Marchinko KB, Arnegard ME, Zhang H, Brady SD, Jones FC, Bell MA, Kingsley DM. 2021. Fitness maps to a large-effect locus in introduced stickleback populations. Proc. Natl. Acad. Sci. U. S. A.. 118:e1914889118.

Schluter D, McPhail JD. 1992. Ecological character displacement and speciation in sticklebacks. Am. Nat.. 140:85–108.

Seppey M, Manni M, Zdobnov EM. 2019. BUSCO: Assessing genome assembly and annotation completeness. Methods Mol. Biol.. 1962:227–245.

Sherman RM, Forman J, Antonescu V, Puiu D, Daya M, Rafaels N, Boorgula MP, Chavan S, Vergara C, Ortega VE et al. 2019. Assembly of a pan-genome from deep sequencing of 910 humans of african descent. Nat. Genet.. 51:30–35.

Simão FA, Waterhouse RM, Ioannidis P, Kriventseva EV, Zdobnov EM. 2015. BUSCO: assessing genome assembly and annotation completeness with single-copy orthologs. Bioinformatics. 31:3210–3212.

Storer J, Hubley R, Rosen J, Wheeler TJ, Smit AF. 2021. The dfam community resource of transposable element families, sequence models, and genome annotations. Mob. DNA. 12:2.

Terekhanova NV, Logacheva MD, Penin AA, Neretina TV, Barmintseva AE, Bazykin GA, Kondrashov AS, Mugue NS. 2014. Fast evolution from precast bricks: genomics of young freshwater populations of threespine stickleback gasterosteus aculeatus. PLoS Genet.. 10:e1004696.

Thompson AC, Capellini TD, Guenther CA, Chan YF, Infante CR, Menke DB, Kingsley DM. 2018. A novel enhancer near the pitx1 gene influences development and evolution of pelvic appendages in vertebrates. Elife. 7.

Urton JR, McCann SR, Peichel CL. 2011. Karyotype differentiation between two stickleback species (gasterosteidae). Cytogenet. Genome Res.. 135:150–159.

Venu V, Harjunmaa E, Dreau A, Brady S, Absher D, Kingsley DM, Jones FC. 2024. Fine-scale contemporary recombination variation and its fitness consequences in adaptively diverging stickleback fish. Nat. Ecol. Evol.. 8:1337–1352.

Wucherpfennig JI, Howes TR, Au JN, Au EH, Roberts Kingman GA, Brady SD, Herbert AL, Reimchen TE, Bell MA, Lowe CB et al. 2022. Evolution of stickleback spines through independent cis-regulatory changes at HOXDB. Nat. Ecol. Evol.. 6:1537–1552.

Xie KT, Wang G, Thompson AC, Wucherpfennig JI, Reimchen TE, MacColl ADC, Schluter D, Bell MA, Vasquez KM, Kingsley DM. 2019. DNA fragility in the parallel evolution of pelvic reduction in stickleback fish. Science. 363:81–84.

Yang C, Zhou Y, Song Y, Wu D, Zeng Y, Nie L, Liu P, Zhang S, Chen G, Xu J et al. 2023. The complete and fully-phased diploid genome of a male han chinese. Cell Res.. 33:745–761.

